# Sustained poly(ADP-ribosyl)ation limits RPA loading and attenuates ATR-CHK1 signaling upon replication fork collapse

**DOI:** 10.1101/2025.10.22.683893

**Authors:** Alexandra Mihuț, Debanjan Ghosh, Dávid Szüts, Roberta Fajka-Boja, Gyula Timinszky

**Affiliations:** Laboratory of DNA Damage and Nuclear Dynamics, Institute of Genetics, HUN-REN Biological Research Centre, Szeged, Hungary; Doctoral School of Multidisciplinary Medical Sciences, University of Szeged, Szeged, Hungary; Doctoral School of Biology, University of Szeged, Szeged, Hungary; Institute of Molecular Life Sciences, HUN-REN Research Centre for Natural Sciences, Budapest, Hungary; Department of Immunology, Albert Szent-Györgyi Medical School, Faculty of Science and Informatics, University of Szeged, Szeged, Hungary

**Keywords:** PARG inhibition, PARylation, ATR-checkpoint, RPA, Replication stress

## Abstract

Poly(ADP-ribosyl)ation (PARylation) is a transient post-translational modification catalyzed by PARP enzymes and reversed by the glycohydrolase PARG. Inhibition of PARG causes sustained PARylation, a toxic state that could be exploited for cancer therapy. However, the mechanisms by which excessive PARylation drives toxicity and cell death remain incompletely understood. Here, we examined how persistent PARylation influences cellular responses to replication stress and DNA damage. We found that during replication stress, the decline of ATR checkpoint activity marks the transition from fork stalling to fork collapse, a context in which stabilized poly(ADP-ribose) reduces chromatin-bound RPA. This loss of RPA, a key activator of the checkpoint, further reduces ATR signaling, revealing a feedback loop between unrestrained PARylation and checkpoint control. Our results identify a checkpoint-dependent consequence of unregulated PARP activity during fork collapse and highlight the PARylation-RPA axis as a critical determinant of PARG inhibitor cytotoxicity.

## Introduction

Replication stress results from a fork slowing or stalling on account of encountering obstacles, dNTP shortage or polymerase inhibition (Berti *et al*, 2020). Mechanistically, resection from double-strand breaks (DSBs) at the fork or helicase-polymerase uncoupling arising from the different regulation of the proofreading polymerase and helicase, creates stretches of single stranded DNA (ssDNA) that are coated by Replication protein A (RPA) (Zhao *et al*, 2020; Byun *et al*, 2005) as a means of protection from nuclease attack and the formation of DNA secondary structures. RPA-coated ssDNA is a signaling platform best known for activating the main replication stress kinase, Ataxia-telangiectasia and Rad3-related (ATR), that stabilizes the forks and drives the checkpoint response to allow DNA replication to resume once the stress has been handled (Saldivar *et al*, 2017; Cimprich & Cortez, 2008).

ATR checkpoint failure during replication stress leads to fork collapse. Concomitantly, RPA exhaustion due to uncontrolled origin firing, deregulated fork remodeling, fork component disintegration, to name a few, will generate unrepaired DNA breaks at the fork and collapse it (Toledo *et al*, 2013; Cortez, 2015; Dungrawala *et al*, 2015), which ultimately can lead to replication catastrophe. In such cases, replication can resume through activation of a nearby dormant origin or via fork restart mechanisms that stabilize and reactivate the stalled fork. (Ge *et al*, 2007; Ibarra *et al*, 2008; Cortez, 2015).

At collapsed or processed forks, DNA ends as generated by DNA damage and DNA processing during replication stress or replication catastrophe are effective substrates for nuclear PARP activation (Rouleau-Turcotte *et al*, 2022; Ortega *et al*, 2025). PARP activation triggers ADP-ribosylation (ADPr) signaling, whose main outcomes include chromatin remodeling, factor recruitment, protein and nucleic acid modification, and ubiquitin-mediated degradation and signaling (Dasovich & Leung, 2023). ADPr is a reversible post-translational modification with roles in the DNA damage response, replication, transcriptional regulation, and cell fate decisions. Poly(ADP-ribose) glycohydrolase (PARG) is the primary enzyme that degrades poly(ADP-ribose) (PAR) chains via its endo- and exo-glycohydrolase activity, thereby terminating ADPr signaling (Crawford *et al*, 2018; Rack *et al*, 2021; Duma & Ahel, 2023).

PARP1 and ADPr have been shown to regulate stalled fork processing such as fork remodeling and fork reversal (Yang *et al*, 2004; Bryant *et al*, 2009a; Berti *et al*, 2013). Clinically, understanding how ADPr shapes fork fate is important because while PARP inhibitors (PARPi) are effective on homologous recombination–deficient backgrounds, PARG inhibitors (PARGi), by leading to persistent poly(ADP-ribosyl)ation (PARylation), can exacerbate replication defects (Slade, 2020). Prior work implicated PARG in fork stability in both challenged and unchallenged cells (Illuzzi *et al*, 2014; Ray Chaudhuri *et al*, 2015) and suggested that PARG inhibition blocks fork restart leading to replication catastrophe, providing a rationale for PARGi as a synthetic-lethal approach in cancers with replication defects (Pillay *et al*, 2019). Nevertheless, the mechanisms by which stabilized PAR chains become toxic remain incompletely understood. Initial models proposed that reduced expression of replication factors sensitizes cells to PARGi-mediated replication catastrophe and subsequent cell death (Pillay *et al*, 2019; Coulson-Gilmer *et al*, 2021). However, recent studies failed to consistently detect low expression of replication genes as an explanation for PARGi-induced replication catastrophe (Coulson-Gilmer *et al*, 2021). Moreover, not all instances of PARGi-mediated cytotoxicity were accompanied by canonical markers of replication catastrophe (Coulson-Gilmer *et al*, 2021). Conflicting findings have also been reported regarding RPA regulation: in some contexts, PARGi reduces RPA, leaving ssDNA unprotected (Illuzzi *et al*, 2014), whereas in others PARGi increases RPA foci, marking impending replication catastrophe (Pillay *et al*, 2019; Coulson-Gilmer *et al*, 2021).

These discrepancies indicate that the precise molecular determinants of RPA reduction upon PAR accumulation remain unresolved. In this study, we addressed the relationship between sustained PARylation and RPA regulation at stressed forks using a variety of replication stressors and cell line models.

## Results

### Sustained PARylation interferes with the replication stress response

PARP1 binds to and is activated at stalled replication forks, and its activity is required to resume replication elongation at stalled forks (Bryant *et al*, 2009b). However, the generated PAR chains are short lived and hardly detectable unless PARG is inhibited or knocked down (Illuzzi *et al*, 2014). To examine the accumulation of PARylation upon replication stress we added PARGi to P19 cells treated with hydroxyurea (HU) or camptothecin (CPT) for various time points. HU blocks the DNA synthesis by inhibiting the ribonucleotide reductase enzyme and thereby decreasing the dNTP pool (Musiałek & Rybaczek, 2021), while CPT binds to the Topoisomerase I–DNA complex, trapping the enzyme on the DNA and obstructing fork progression (Li *et al*, 2017). As expected, the signal of PARylation increased over time, which was only detectable in the presence of PARGi (Fig. 1A). However, the accompanying signaling events were different between HU and CPT treatment. As short as 1 h incubation elicited only mild replication stress in CPT-treated cells indicated by very low levels of phosphorylated CHK1, RPA32 and H2AX. These signaling events increased over time, associated with CHK2 phosphorylation and high levels of γH2AX at 6 h, denoting ataxia-telangiectasia mutated (ATM) activation and DNA damage response, respectively (Zannini *et al*, 2014) (Fig. 1A). In contrast, HU treatment induced excessive amount of phospho-CHK1 both at S317 and S345 sites, known to be primarily phosphorylated by ATR (Zhao & Piwnica-Worms, 2001), already at 1 h of incubation, the shortest treatment time we assessed, and decreased over time. The γH2AX was constantly high at all of the time points examined, while CHK2 phosphorylation was barely detectable. In spite of decreasing pCHK1, the phosphorylated form of RPA32 at T21 and S4/S8 sites increased over time upon HU treatment, unless combined with PARGi. PARGi co-treatment resulted in the marked reduction in the level of phosphorylated RPA32 in the case of HU but was less prominent upon CPT treatment at the examined time-points. In addition, combination of PARGi with HU further decreased CHK1 phosphorylation while keeping the level of γH2AX unchanged, suggesting that sustained PARylation interferes with replication stress signaling rather than with the DNA damage response (Fig. 1A).

**Figure 1.**
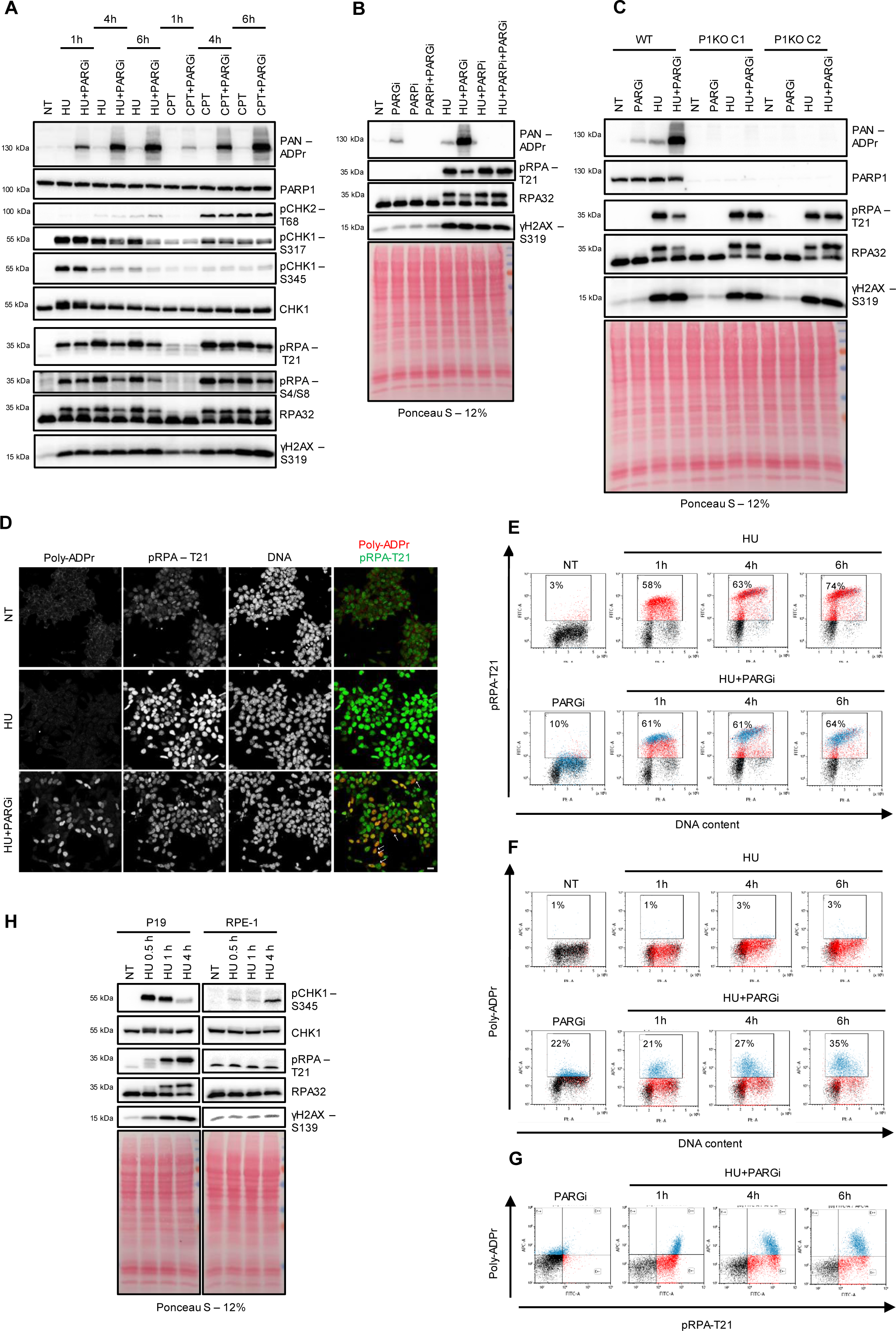
Replication stress response in the presence of persistent PARylation. (**A**) P19 cells were treated with HU (0.5 mM) or CPT (5 µM), with or without PARGi (5 µM), for the indicated time points, lysed and analyzed by western blotting with the indicated antibodies. (**B**) P19 cells were treated with PARGi (5 µM), PARPi (5 µM), HU (0.5 mM) and all combinations thereof for 4 hours then lysed and analyzed by western blotting with the indicated antibodies. Ponceau S staining was used as a loading control. (**C**) P19 WT and PARP1 KO cells (clones P1KO C1 and P1KO C2) were treated with PARGi (5 µM), HU (0.5 mM) and their combination for 7 hours then lysed and samples were analyzed by western blotting with the indicated antibodies. Ponceau S staining was used as a loading control. (**D**) P19 cells were treated with HU (0.5 mM) with or without PARGi (5 µM) for 4 hours, fixed and analyzed by immunofluorescence with the indicated antibodies. Scale bar: 20 µm. Arrows indicate cells with high PAR and low pRPA levels. (**E, F, G**) P19 cells were treated with HU (0.5 mM) with or without PARGi (5 µM) for the indicated time points, fixed and analyzed by FACS with the indicated antibodies. The pRPA-T21^+^ cells are colored in red and PAR^+^ cells are blue throughout all of the FACS images. (**H**) P19 and RPE-1 cells were treated with HU (0.5 mM for P19, 1 mM for RPE-1) for the indicated time points, lysed and analyzed with the specified antibodies.

To confirm the involvement of PARP activity in the cellular response to HU-induced replication stress, we co-treated the cells with the PARPi, Olaparib. PARPi completely blocked PARylation, but left the RPA32 and H2AX phosphorylation unaltered during the 4 h co-incubation with HU and completely abrogated the PARGi-induced reduction in RPA32 phosphorylation (Fig. 1B). Note that neither PARGi nor PARPi triggered pRPA or γH2AX without HU-treatment, indicating that perturbing PARylation during regular replication does not induce replication stress or DNA damage. The combined treatment with HU, PARPi and PARGi revealed that the excess PAR signal was responsible for the reduced level of phospho-RPA32 and allowed us to exclude that this would arise from an off-target effect of the PARG inhibitor. In addition, we established PARP1 knock-out (KO) derivatives of P19 cells using the CRISPR-Cas9 nickase gene-editing approach. Similar to PARPi treatment, the replication stress and DNA damage response were not different in PARP1 KO cells as compared to wild-type cells, but PARGi co-treatment failed to reduce the phosphorylation of RPA32 in the absence of PARP1 (Fig. 1C). Altogether, the PARP1 KO and PARP inhibition experiments indicate that the phosphorylation status of RPA32 during HU-treatment depends on the presence and activity of PARP1.

We examined the reciprocal correlation between PARylation and pRPA-T21 by immunofluorescence (Fig. 1D). Adding PARGi decreased the intensity of pRPA-T21 staining in HU-treated samples, especially in those with high PARylation signal (Fig. 1D, arrows). We also followed the changes of these signals with flow cytometry over several time-points. HU treatment alone induced RPA32 phosphorylation in S-phase cells from 1 h of incubation, with the signal increasing over time (Fig. 1E). When PARGi was added, the accumulated PARylation signal could be detected, which was the highest in early S-phase (Fig. 1F). Notably, the pRPA-T21 fluorescence intensity decreased in PAR-positive cells from 4 h of HU and PARGi co-treatment (Fig. 1G), which was most evident in early S-phase cells (Fig. 1E, lower row). It has been shown previously, that the HU-induced hyperphosphorylation of RPA32 was impaired in PARG knockdown HeLa cells upon longer treatment, i.e. over 12 h of incubation (Illuzzi *et al*, 2014). Therefore, we compared the response of P19 cells to HU with that of HeLa and the human retinal pigment epithelium (RPE)-1 cells after short- and long-term incubation (Supplemental Figure 1). We found that extending the HU treatment in P19 cells from 4 to 16 h resulted in a decrease in CHK1 and RPA32 phosphorylation, accompanied by an increase in γH2AX and PARylation signals. CHK1 and RPA32 phosphorylation were both reduced under sustained PARylation. In contrast, the CHK1 phosphorylation slightly increased over time whereas the phospho-RPA signal was constant throughout the treatment in HeLa cells. The RPE-1 cell line followed a similar trend to HeLa, opposite to P19, with the checkpoint increasing with treatment time rather than declining. The γH2AX and robust PARylation signals were not induced by HU in the RPE-1 cells. In HeLa cells, PARylation, but not γH2AX, was induced after prolonged treatment however it failed to reduce CHK1 or RPA phosphorylation. This suggests that the presence of PARylation and RPA is necessary but not sufficient for them to correlate and that a third component is required. We continued to compare the kinetics of early signaling events in P19 versus RPE-1 cells. While CHK1 became phosphorylated rapidly upon HU-treatment and declined with incubation time, pRPA-T21 and γH2AX gradually increased in P19 cells (Fig. 1H). In contrast, we could detect only a modest pCHK1 level in RPE-1 cells, accumulating from 4 h of treatment, and no increase in pRPA-T21 and γH2AX signal (Fig. 1H).

These results identify the P19 cell to be particularly sensitive to HU-induced fork stalling even upon short treatment, resulting in fork collapse, recorded by strong CHK1 and RPA32 phosphorylation accompanied by DNA damage response markers γH2AX and PAR early on. Preserving PARylation by blocking PARG activity reveals that it interferes with stress signaling by reducing the CHK1 and RPA32 phosphorylation, but does not alter the DNA damage signaling, as it leaves γH2AX signal unchanged.

### Inhibiting checkpoint signaling at stalled replication forks elevates PARylation levels and reduces RPA32 phosphorylation

We followed the nucleotide incorporation upon HU treatment by adding the nucleotide analogue 5-bromo-2′-deoxyuridine (BrdU), and we found that it markedly decreased after 30 minutes and was completely blocked at 1 h in P19 cells (Supplemental Figure 2). In contrast, the DNA replication continued at a low frequency in the presence of HU in RPE-1 within the same timeframe (Supplemental Figure 2), mirroring the differences in the early signaling events in these two cell lines. We assumed that the differences of replication stress signals, like CHK1 and RPA32 phosphorylation evoked by HU and CPT also depended on the progression of DNA replication. While HU completely blocked nucleotide incorporation as early as 1 h treatment in P19, CPT only decreased it, slowing the cells’ propagation to the late S-phase in P19, resulting in accumulation of the cells in early S-phase after 4h incubation (Fig. 2A). Additionally, we tested two other drugs, Aphidicolin (APH) and Etoposide (ETO), which also stalled or slowed down the replication fork, respectively (Fig. 2A). APH induced similar response to HU in terms of PARylation, CHK1, RPA32 and H2AX phosphorylation at 4h of incubation (Fig. 2B). The combination of APH with PARGi reduced the level of phosphorylated CHK1 and RPA32, but not γH2AX. In contrast, ETO and CPT induced higher pRPA and γH2AX level but lower pCHK1 signal suggesting that DNA damage rather than replication stress was the underlying mechanism. Interestingly, while both ETO and CPT reduced CHK1 phosphorylation when combined with PARGi, they did not markedly decrease the pRPA signal (Fig. 2B).

**Figure 2.**
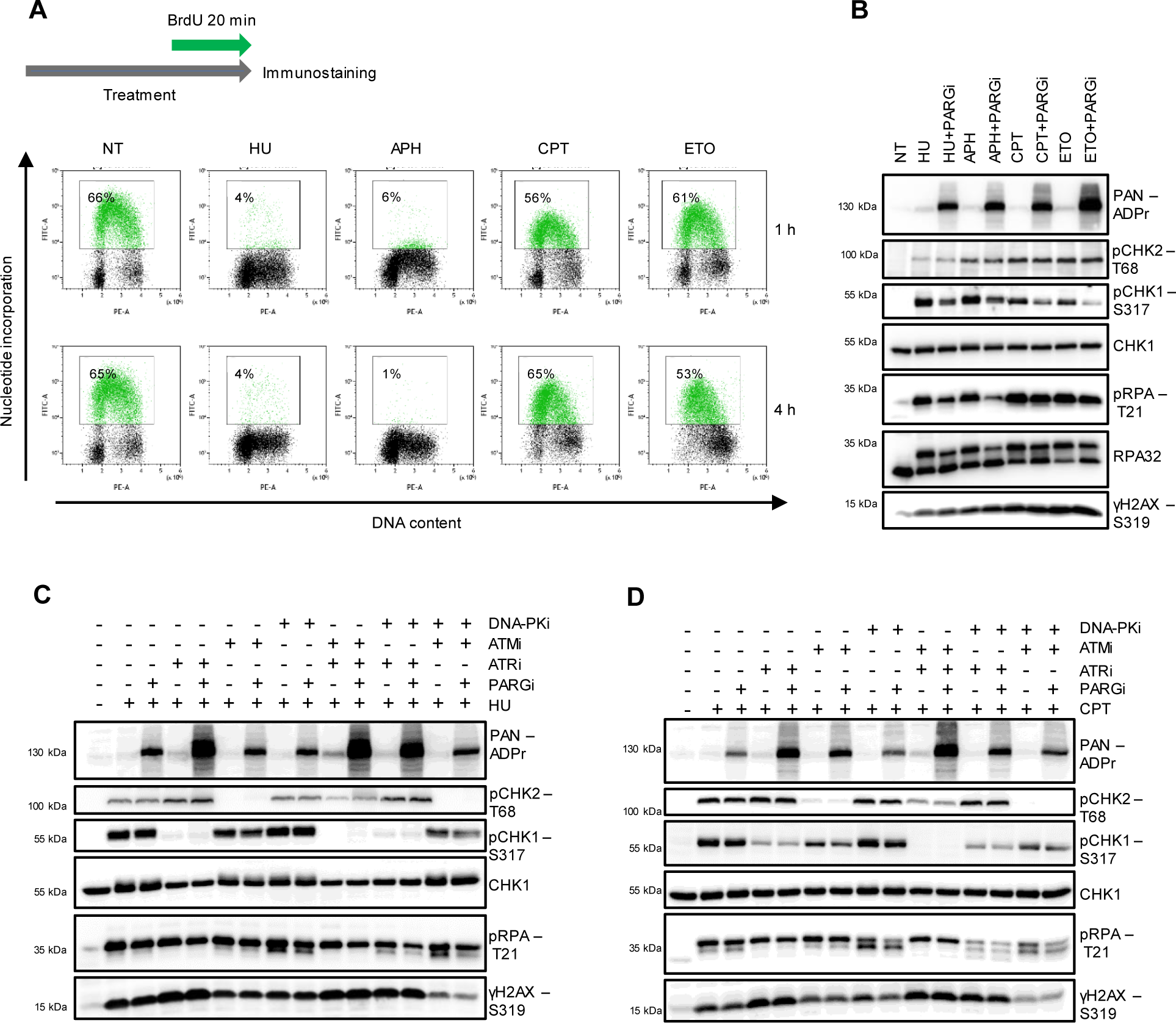
Differences in HU and CPT-induced early replication stress signaling. (**A**) P19 cells were treated with HU (0.5 mM), APH (10 µM), CPT (5 µM) and ETO (10 µM) for 1 h. BrdU was added for the last 20 minutes to show the nucleotide incorporation. The cells were then fixed and stained with anti-BrdU and propidium iodide (DNA content), and then the samples were analyzed with flow cytometry. FACS images of a representative experiment are shown. (**B**) P19 cells were treated with HU (0.5 mM), APH (10 µM), CPT (5 µM) and ETO (10 µM) with or without PARGi (5 µM) for the indicated time points. Cells were then lysed and subjected to western blotting with the indicated antibodies. (**C, D**) P19 cells were pre-treated with ATRi (5 µM), ATMi (5 µM) and DNA-PKi (5 µM) for 30 minutes after which HU (0.5 mM) (**C**) or CPT (5 µM) (**D**) were added for an additional 1 hour and 30 minutes with or without PARGi (5 µM). Cells were then lysed and subjected to western blotting with the indicated antibodies.

To further analyze why HU- and CPT-induced PARylation reduced pRPA with different efficiency we looked at the early stress response as controlled by the three main replication stress and DNA damage kinases, ATR, ATM and DNA-dependent protein kinase (DNA-PK). The kinases were inhibited for 30 minutes ahead of the HU and CPT treatments, which were then added to the cells for an additional 1 hour and 30 minutes together with the kinase inhibitors. At this short incubation time both HU and CPT induced RPA32 and H2AX phosphorylation, but the reduction in pRPA level in the presence of PARGi was not significant yet (Fig. 2C and D). However, the difference between HU and CPT checkpoint signaling was already detectable: while HU mainly triggered CHK1 phosphorylation controlled by ATR activity, CPT relied on CHK2 phosphorylation dependent on ATM, a kinase activated at DNA breaks. On the other hand, there was an interplay between these signals: when ATR was inhibited under HU treatment, CHK2 phosphorylation was elevated, in turn, ATM inhibitor (ATMi) moderately decreased the pCHK1 signal (Fig. 2C). The dominance of ATR control upon HU-treatment was also indicated by elevated PARylation and γH2AX when ATR was inhibited and consequential fork collapse. Co-treatment with ATR inhibitor (ATRi) and PARGi resulted in less phospho-RPA32 signal during HU-induced replication stress even at this early incubation time. However, phosphorylation of the T21 residue of RPA32 was dependent on the activity of DNA-PK, as it was previously reported (Liu *et al*, 2012), and the γH2AX signal was decreased by ATM or DNA-PK inhibition or their combination (Fig. 2C and D). In contrast to HU, in the case of CPT-induced replication stress ATRi did not elevate PARylation when DNA-PK was also inhibited, indicating that in this case the signaling was dominated by the latter (Fig. 2D). Nonetheless, the CPT-induced pRPA-T21 signal was stronger decreased when DNA-PK inhibitor (DNA-PKi) was combined with ATRi or ATMi, than with DNA-PKi alone (Fig. 2D).

Taken together, our results are consistent with HU-induced replication fork stalling being governed by the ATR/CHK1 axis while the lesions caused by CPT treatment drive predominantly ATM/CHK2 and DNA-PK activation. Yet, these pathways work in concerted ways, finally leading to the observed stress signals, RPA32 and H2AX phosphorylation. Resolving the replication stress differs in the case of HU from that of CPT with immediate CHK1 and RPA32 phosphorylation, which gradually decrease as PARylation increases suggesting that P19 cannot maintain fork stalling leading to their collapse.

### Suspending the ATR-CHK1 checkpoint control leads to decreased level of chromatin-bound RPA32 in the presence of PARGi

Next, we raised the question if sustained PARylation also affected the amount of chromatin-bound RPA32. We HU-treated P19 cells for 1 and 4 hours with or without PARGi and tested the amount of cytoplasmic, nuclear and chromatin-bound RPA32 (Fig. 3A). The faster migrating band of RPA32, which represent the non-phosphorylated form, was found mostly in the cytoplasmic fraction under all conditions, while the slower migrating phosphorylated band was associated with the chromatin fraction in HU-treated samples even at shorter incubation time. The amount of phospho-RPA32 protein was decreased in the presence of PARGi and showed inverse correlation with the PARylation signal (Fig. 3A). It should be noted that the reduction of HU-induced phospho-RPA32 by PARGi was not accompanied by the increase of its non-phosphorylated form suggesting that the overall level of DNA-bound RPA32 is reduced by sustained PARylation. However, by adding PARPi to the cells we found that PARylation did not mediate the loading of RPA32 on chromatin upon replication stress, but PARPi prevented the effect of PARGi in reducing DNA-bound RPA32 (Fig. 3B), further supporting that poly(ADP-ribosyl)ation was needed for decreasing the binding of RPA32 to DNA. Despite that all tested drugs led to high levels of PARylation, the accompanied reduction of DNA-bound RPA32 was more characteristic to fork-stalling drugs, such as HU and APH, than to the ones inducing DNA breaks like CPT and ETO (Fig. 3C).

**Figure 3.**
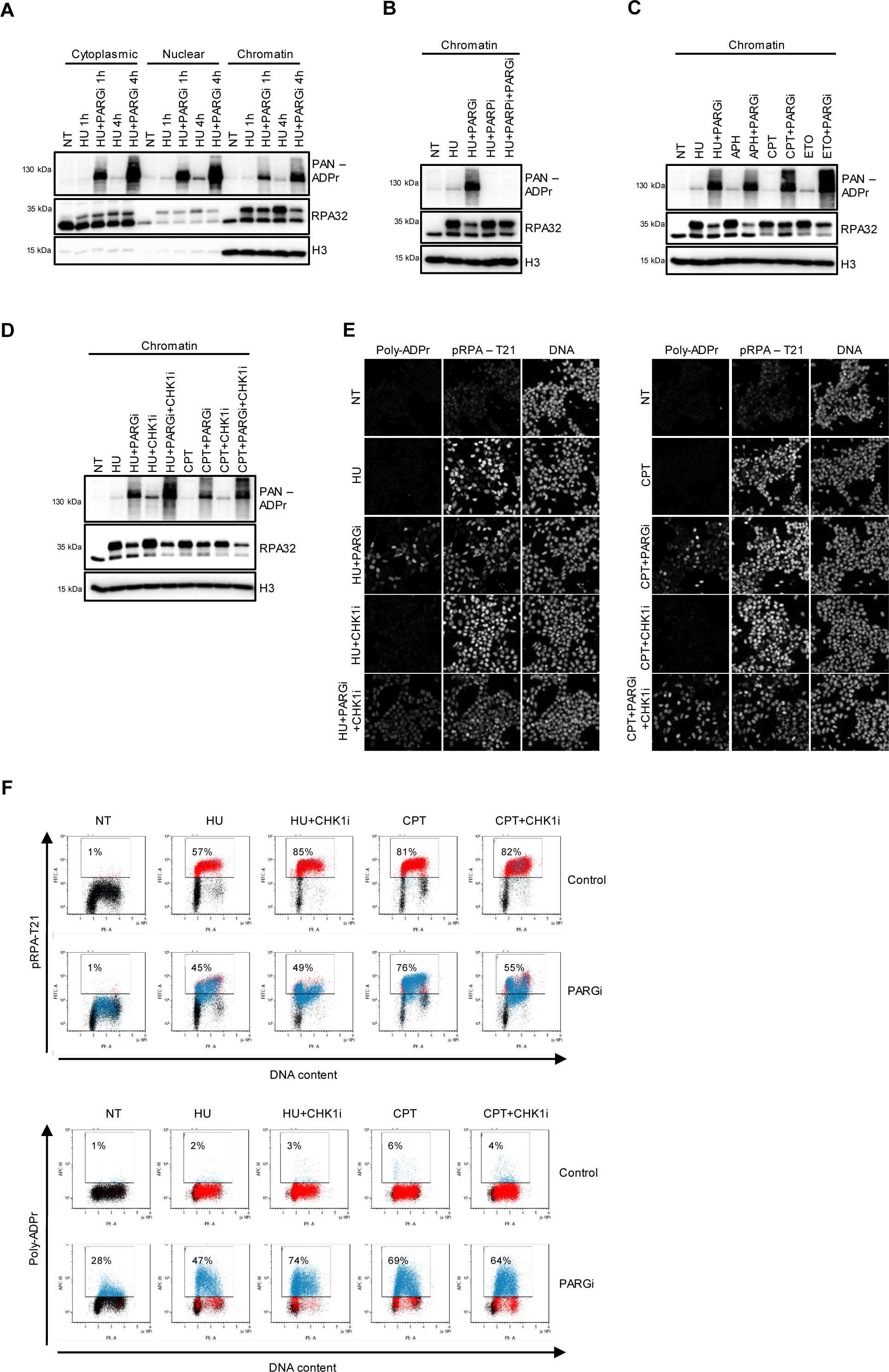
Correlation between loss of the ATR-checkpoint and the level of chromatin-bound RPA32. **(A)** P19 cells were treated with HU (0.5 mM) with or without PARGi (5 µM) for the indicated time points and subjected to cell fractionation to assess the levels of PARylation and RPA32 in cytoplasmic, nuclear or chromatin fraction by western blotting. **(B)** PARylation and RPA32 levels on the chromatin of P19 cells treated with HU (0.5 mM) with or without PARGi (5 µM), PARPi (5 µM) and their combination for 4 hours assessed by western blotting. **(C)** PARylation and RPA32 levels on the chromatin of P19 cells treated with HU (0.5 mM), APH (10 µM), CPT (5 µM) and ETO (10 µM) with or without PARGi (5 µM) for 4 hours assessed by western blotting. **(D)** PARylation and RPA32 levels on the chromatin of P19 cells treated with HU (0.5 mM) and CTP (5 µM) with or without PARGi (5 µM) in the presence or absence of CHK1i (5 µM) for 4 hours assessed by western blotting. (**E, F**) PAR and pRPA levels of P19 cells treated with HU (0.5 mM) and CTP (5 µM) with or without PARGi (5 µM) in the presence or absence of CHK1i (5 µM) for 4 hours assessed by immunofluorescence and FACS.

As we found in our experiments (Fig. 2C), the ATR/CHK1 axis dominated the HU-induced replication stress response. Co-treatment of P19 cells with HU and the CHK1 inhibitor (CHK1i) for 4 h led to increased signals of pRPA-T21, PARylation and γH2AX, while the PARGi-induced reduction of RPA32 phosphorylation was retained (Supplemental Figure 3A). Therefore, we tested if the CHK1i also interfered with RPA32 accumulation on chromatin. CHK1i elevated the amount of PARylation both in the case of HU- and CPT-treatment and the chromatin-bound RPA32 was more efficiently reduced by PARGi when CHK1i was present (Fig. 3D). Importantly, CHK1i also sensitized RPE-1 cells to HU-induced stress: high PARylation, induction of γH2AX, reduced pRPA-T21 levels and less chromatin-bound RPA32 were detected when HU treatment was combined with CHK1i in the presence of PARGi (Supplemental Figure 3B). We further corroborated that this effect was due to checkpoint suppression by ATR inhibition (Supplemental Figure 3C). The distribution of PARylation and pRPA-T21 signal was also followed with fluorescence microscopy and flow cytometry under these conditions in P19 cells (Fig. 3E and F). Both methods showed that CHK1i increased the number of cells with PARylation and promoted the reduction of phospho-RPA32 intensity when added either to HU- or CPT-treated P19 cells.

These results indicate that excessive accumulation of PARylation prevents RPA loading on and/or promotes its dissociation from chromatin upon replication stress, which decreases the ATR/CHK1 activation. Pharmacological inactivation of CHK1 further elevates PAR levels and decreases chromatin-bound RPA, which hints at a PAR-mediated feedback loop to counter replication fork collapse.

### Disruption of replication forks and opening new origins upon HU treatment in P19 cells

In most cells, short HU treatment causes replication fork stalling, which can be restarted; in contrast, long treatment (over 12 hours) leads to fork collapse, when replication is rescued by new origin firing (Petermann *et al*, 2010). PARGi-mediated pRPA reduction was detected only upon longer treatment with HU in the presence of CHK1i in case of RPE-1 cells (Supplemental Figure 3B and C), suggesting that fork collapse rather than fork reversal emerged. Indeed, we found that the reduction in pRPA-T21 level also happened in RPE-1 cells deficient for BRCA1, a homologous recombination factor playing role in fork reversal (Supplemental Figure 4A). In contrast, we detected that the protein level of Timeless (TIM) and Proliferating cell nuclear antigen (PCNA), components of the replication machinery, decreased in chromatin fraction as early as 1 h incubation with HU in P19 cells (Fig. 4A), which indicated fork collapse and the dissociation of the replisome at an unusually early time point. This may be due to the failure to maintain high levels of ATR activity, as we observed that CHK1 phosphorylation was the highest at already 30 minutes and decreased over time (Fig. 1H). Inhibition of the ATR/CHK1 pathway gives rise to fork collapse and the opening of new origins (Cimprich & Cortez, 2008). When we added ATRi to HU-treated samples, it elevated the amount of chromatin-bound TIM and PCNA, suggesting that replication machineries were built up at new origins (Fig. 4A). ATRi also elevated the level of chromatin-bound RPA32 but did not change its PARGi-mediated reduction most evident at 4 h of incubation time (Fig. 4A). We experienced a similar pattern when using CHK1i upon HU treatment (Supplemental Figure 4B), however, the recruitment of TIM to chromatin upon CPT and CHK1i co-treatment was weaker compared to HU (Fig. 4B), again pointing at the differences between the stress response elicited by these drugs. Decrease of TIM and PCNA in chromatin fraction was accompanied with increase of γH2AX in P19 cells (Fig. 4C). These results further support that P19 cells are extremely sensitive to HU treatment, responding with a very early onset of fork collapse. In contrast, in RPE-1 cells, the levels of TIM and PCNA did not decrease on chromatin and there was no γH2AX signal upon 4 h HU treatment (Fig. 4C), suggesting that there was no fork collapse during this time.

**Figure 4.**
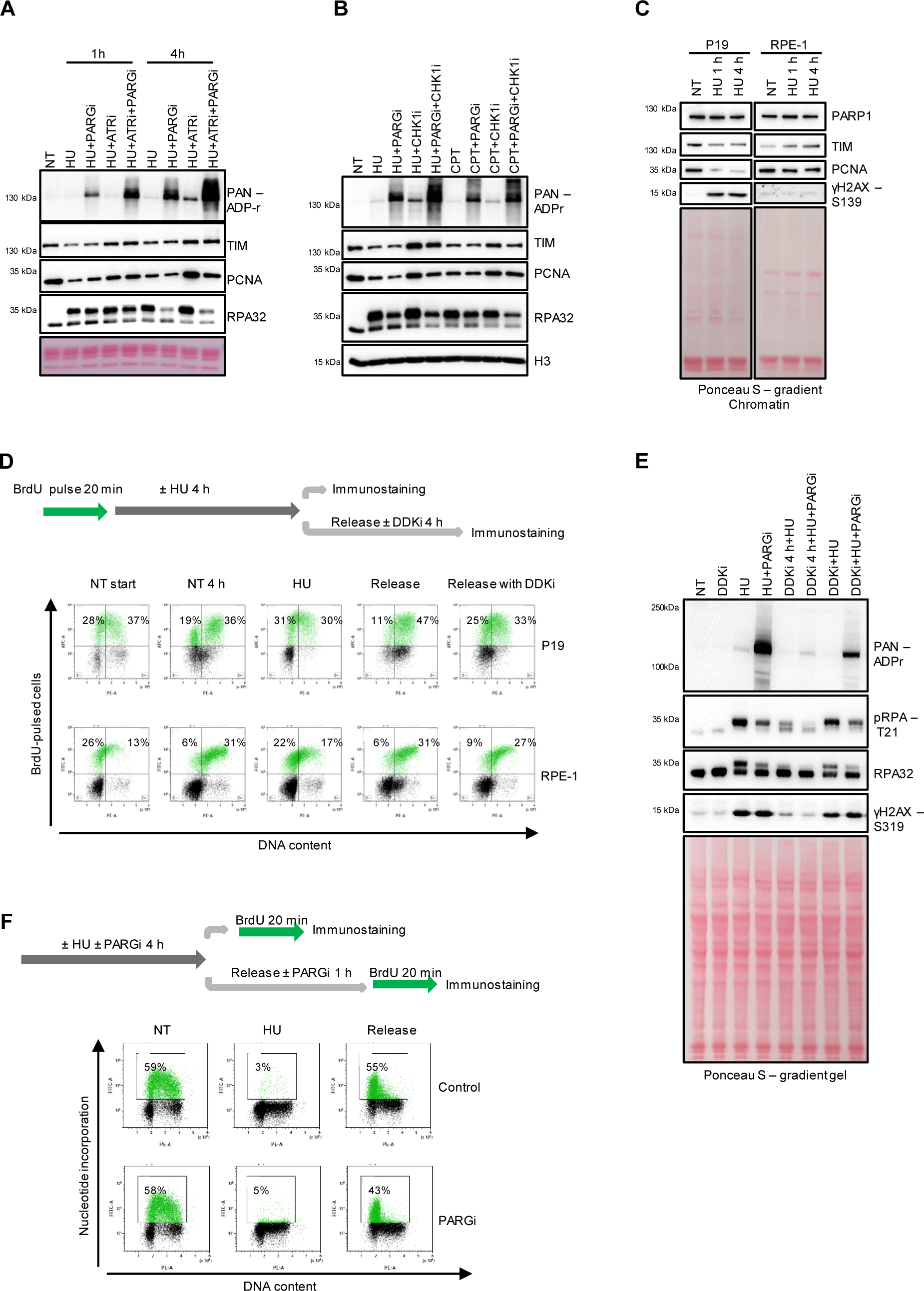
Detection of HU-induced fork collapse and new origin firing. **(A)** P19 cells were treated with HU (0.5 mM) with or without PARGi (5 µM) in the presence or absence of ATRi (5µM) for 1 or 4 h and subjected to chromatin fractionation. PARylation, TIM, PCNA and RPA32 levels were assessed by western blotting. **(B)** P19 cells were treated with HU (0.5 mM) or CTP (5 µM) with or without PARGi (5 µM) in the presence or absence of CHK1i (5 µM) for 4 hours and PARylation, TIM, PCNA and RPA32 levels were assessed by western blotting. **(C)** P19 and RPE-1 cells were treated with HU (0.5 mM for P19, 1 mM for RPE-1) for the indicated time points. Chromatin fractionation and western blotting was performed to assess fork components level together with DNA damage as specified by the antibodies. **(D)** P19 and RPE-1 cells were incubated with BrdU (NT start) to pulse-label the S-phase cells and then treated with HU (0.5 mM) for 4 h in the presence or absence of DDKi (1 µM), or left untreated. The cells were then fixed or released for 4 h. The propagation of labelled cells throughout the cell cycle was followed by anti-BrdU and propidium iodide (DNA content) staining, and analyzed with flow cytometry. FACS images of a representative experiment are shown. **(E)** P19 cells were treated with HU (0.5 mM) with or without PARGi (5 µM) for 4 hours. DDKi (1 µM) was either pre-incubated for 4 hours (lanes 5 and 6) to the HU or added at the same time (lanes 7 and 8). Cells were lysed and the samples were subjected to western blotting by the indicated antibodies. **(F)** P19 cells were treated with HU (0.5 mM) with or without PARGi (5 µM) for 4 hours. The cells were then fixed or released for 1 h. BrdU was added for the last 20 minutes. The samples were stained with anti-BrdU (nucleotide incorporation) and propidium iodide (DNA content), and then the samples were analyzed with flow cytometry. FACS images of a representative experiment are shown.

Our results so far suggest that HU-induced replication stress rapidly leads to fork collapse in P19 cells, which is normally controlled by the ATRi/CHK1 pathway. As we observed that the phosphorylation of the CHK1 decreased over time upon HU-treatment in P19 (Fig. 1A), which should allow the firing of new origins. To test this hypothesis, we pulse labeled S-phase cells with BrdU and then followed cells along the cell cycle by determining their DNA content with propidium-iodide staining and flow cytometry. BrdU-labelled P19 cells reached late S/G2-phase within 4 h and a proportion of cells already divided and was in G1 of the next cell cycle (Fig. 4D). In contrast, HU-treated cells could not replicate and increase their DNA content unless they were released from the treatment. Cell Division Cycle 7 (Cdc7), also known as Dbf4-dependent kinase (DDK) is an enzyme responsible for the activation of the replicative helicase MCM and for the initiation of DNA replication at multiple origins (Fragkos *et al*, 2015). In our experiments, DDK inhibitor (DDKi) abolished the DNA replication in released cells, proving that new origin firing was indeed required for DNA replication after HU-treatment (Fig. 4D). This was not the case for RPE-1 cells, as their progress along cell cycle was not hindered by DDKi after release from the 4 h HU-block (Fig. 4D), indicating that their forks were stalled but not collapsed.

Pre-incubation of P19 cells with DDKi prior to HU treatment would restrain them from entering S-phase and starting DNA replication. These cells were insensitive to HU-induced stress, showing very low level of PARylation, pRPA-T21 and γH2AX (Fig. 4E). The simultaneous treatment of DDKi with HU, where only the ongoing forks were affected by the burden, resulted in a moderate cell response. These results indicate that the signals detected upon replication stress of P19 stemmed from both stalled replication forks and the newly opened origins. To follow the DNA replication from the new origins after replication stress and fork collapse, we released the cells after 4 h HU-treatment and monitored the nucleotide incorporation by adding BrdU for the last 20 min of the 1 h incubation time. Nucleotide incorporation was reduced by PARGi-treatment after release from HU-block, suggesting that the sustained PARylation inhibits the opening of new origins or slows DNA replication (Fig. 4F).

Altogether these experiments reveal that P19 cells respond with enhanced sensitivity to HU-triggered replication stress. Their immediate CHK1 phosphorylation highlights a strong checkpoint control followed by rapid fork collapse and opening of new origins as a salvage pathway. The pRPA-T21 and γH2AX together with high PARylation suggest the initiation of DNA repair at the collapsed forks.

## Discussion

We addressed the effect of sustained poly(ADP-ribosyl)ation by PARG inhibition on the replication stress response focusing on its early and acute effects. Our data support a model in which replication fork collapse activates PARP1 and drives PAR accumulation; stabilized PAR chains then reduce chromatin-bound RPA (including phosphorylated isoforms) and decrease ATR–CHK1 signaling. This causal direction reconciles why PAR and low RPA coincide with γH2AX a proxy for DSB-associated ATM and DNA-PK signaling upon PARG inhibition during replication stress, and why conditions that prevent collapse do not show RPA reduction. While it has been reported that chronic RNAi-mediated PARG depletion increases γH2AX signal (Illuzzi *et al*, 2014), during both short- and long-term PARG inhibition, sustained PARylation does not alter the fork collapse-induced γH2AX. Thus, PARylation does not appear to act through the modulation of ATM or DNA-PK activity during replication stress.

Different replication stressors and DNA damage inducing agents reveal that PARylation levels do not always correlate with the amount of RPA reduction. HU and aphidicolin rapidly stall forks – no BrdU incorporation is detectable within one hour – yet they are not the strongest inducers of PARylation; nevertheless, they produce the most pronounced reduction in chromatin-bound RPA and CHK1 phosphorylation. By contrast, camptothecin and etoposide impose delayed inhibition of DNA synthesis, consistent with fork slowing rather than outright stalling. Camptothecin induces fork reversal (Ray Chaudhuri *et al*, 2012), which limits ssDNA accumulation and accounts for the comparatively modest chromatin-bound RPA and pCHK1 under these conditions. Thus, the functional link between PARylation and RPA regulation manifests primarily upon fork-stalling.

Checkpoint status dictates whether fork stalling transitions to fork collapse. In P19 embryonic cells, HU alone rapidly triggers fork collapse: we observe loss of fork components TIM and PCNA from chromatin and γH2AX induction, hallmarks of fork collapse. Under these conditions, sustained PARylation reduces pRPA and pCHK1 in P19, whereas HeLa cells do not show the same response to HU. In non-tumorigenic RPE-1 cells, HU elicits only a modest response – RPA phosphorylation appears after prolonged exposure without overt DNA damage markers. However, combining HU with checkpoint inhibition in RPE-1 provokes hyperphosphorylated RPA species, γH2AX, and when PARGi is present it leads to elevated PARylation and the reduction of hyperphosphorylated RPA forms and that of chromatin-bound RPA. Inhibiting the checkpoint during CPT exposure – otherwise a weak inducer of PARylation-linked pRPA reduction – likewise produces a robust drop in chromatin-bound RPA and pRPA. Collectively, replication stress combined with checkpoint loss promotes fork collapse, which activates PARP1; the resultant persistence of PAR chains then limits RPA loading onto chromatin.

Yet, the way sustained PARylation lowers chromatin-bound RPA remains elusive. RPA has high affinity for single-stranded DNA. One could envision that the reduced amount of chromatin-bound RPA was due to the inability of RPA to efficiently bind ssDNA (or due to its increased dissociation from ssDNA) through its ADP-ribosylation. A similar scenario would be if RPA got depleted because of excessive ssDNA formation, the ssDNA not bound by RPA would be attacked by nucleases leading to DNA lesions and ultimately to replication catastrophe (Toledo *et al*, 2013). The excessive increase in DNA lesions is associated with an increased γH2AX signal, something we do not observe at least up to 24 hours of PARGi treatment, therefore, it is rather unlikely that single-stranded DNA would be left “naked” through ADP-ribosylation-mediated sequestration or depletion of RPA.

Further assuming that the amount of ssDNA remains constant upon PARG inhibition, RPA reduction could be achieved through its replacement by Rad51, which is facilitated by BRCA1. However, it is unlikely to be the case, because in our experiments PARGi leads to RPA reduction in BRCA1 deficient cells too. Alternatively, the reduction of RPA on ssDNA could be due to the PARylation of DNA (Groslambert *et al*, 2021; Talhaoui *et al*, 2016). While there is no experimental evidence for the ADP-ribosylation of DNA during replication stress, the ADP-ribosylation of DNA can be achieved by a bacterial toxin, DarT, which leads to massive replication stress and increased RPA phosphorylation – rather than the reduction in chromatin-bound RPA we observe upon PARGi – in cells lacking TARG1, an ADP-ribosyl hydrolase critical to remove the DarT generated ADP-ribose from DNA (Tromans-Coia *et al*, 2021). PARP1 was shown to ADP-ribosylate the 3’ terminal phosphate DNA ends, at least, in vitro (Matta *et al*, 2020). Even if this happened during replication stress, it is rather unlikely that the ADP-ribosylation of DNA ends could alter RPA binding along ssDNA strands. Thus DNA ADP-ribosylation is not a likely mechanism that could trigger the reduction of RPA on ssDNA.

On the other hand, PARP1 activity and PARylation were reported to promote the accumulation of reversed forks and slows fork restart by PAR-dependent inhibition of RECQ1. While, fork reversal is primarily catalysed by RAD51 and the fork-remodelling translocases, such as HLTF, SMARCAL1, ZRANB3, PARylation appears to impact fork reversal largely through its effect on RECQ1 (Zellweger *et al*, 2015). We have not measured fork reversal and fork restart specifically, but the BrdU incorporation assays showed that PARGi reduced BrdU incorporation upon the release of cells from HU treatment. Both decreased fork speed and decreased origin firing might reduce BrdU incorporation. While PARylation hasn’t been implicated in origin firing, the observed reduced BrdU incorporation could be due to the above-mentioned effects of PARylation on fork speed through decreasing RECQ1 activity. Sustained PARylation upon PARGi treatment should thus reduce fork speed and promote fork reversal, which during HU treatment could lower the amount of ssDNA.

In our experimental conditions, the RPA-reducing effect of PARGi was only evident during fork collapse when the DNA lesions lead to efficient PARP1 activation. The proteome analysis of replisome dynamics in response to fork stalling by HU revealed that PARG levels at the replisome get reduced shortly after HU treatment, which is accompanied by the enrichment of macroH2A histones, that can bind ADP-ribose (Dungrawala *et al*, 2015). These results suggest that replication stress induced by HU leads to increased ADP-ribosylation. Considering that the cellular responses to replication stress are to counter replication stress, ADP-ribosylation might also be considered as a safeguarding mechanism that can slow fork speed and that way reduce replication stress or may delay the onset of replication catastrophe, were it merely due to excess ssDNA over RPA. However, when PARylation is deregulated though PARG inhibition, persisting PARylation may interfere with efficient fork restart and recovery from replication stress that may underlie its clinically relevant effect being synthetic lethal with the loss of a number of genes implicated in the replication stress response (Pillay *et al*, 2019).

## Materials and methods

### Cell lines and culture conditions

The P19 embryonic cell line was a kind gift from Professor István Raskó (Rasko *et al*, 1993). PARP1 knockout (KO) P19 cells were generated in-house using CRISPR-Cas9 nickase-mediated genome editing. RPE-1 *TP53^-/^*^-^ and *TP53^-/-^ BRCA1^-/-^* cells were generously provided by Alan D. D’Andrea (Lim *et al*, 2018). The HeLa cell line was authenticated through STR profiling (Eurofins Genomics), showing a 100% match with HeLa (amelogenin + 12 loci) according to the Cellosaurus database (Bairoch, 2018).

RPE-1 cells were cultured in DMEM/F-12, while P19 and HeLa cells were cultured in high-glucose DMEM. Culture media were supplemented with 10% FBS, 100 U/ml penicillin and 100 μg/ml streptomycin. For HeLa cells, the medium was additionally supplemented with non-essential amino acids. Cells were maintained in a humidified incubator at 37 °C with 5% CO_2_.

### CRISPR-Cas9 nickase-mediated knockout generation

We generated PARP1 KOs in the P19 cell line by means of CRISPR mediated genetic editing. We used a modified version of the CRISPR-Cas9 called D10A or nickase, pSpCas9n(BB)-2A-Puro (PX462) V2.0, which was a kind gift from Feng Zhang lab (Addgene, plasmid #62987) (Ran *et al*, 2013). The 4 guide-RNAs (gRNA) were cloned into pUC-H1-gRNA, a kind gift from Shenglin Huang lab (Addgene, plasmid #61089) (Zheng *et al*, 2014). DharmaFECT was used for the plasmid transfection according to the manufacturer’s instructions. The KOs were validated by PCR and Western blotting. The following gRNAs were used to generate a 1 kb deletion spanning over exon 6 and 7 of the mouse PARP1 gene:

gRNA1: 5’ – GGTGCCGTCAGGAGAGTCAG – 3’

gRNA2: 5’ – CAGCTCCTTCAGGTCGTTGG – 3’

gRNA3: 5’ – CGACGCTTATTACTGTACTG – 3’

gRNA4: 5’ – TGGCCTGAACACTCCTTGCA – 3’

### Western blotting

Cells were treated for the indicated time points in fresh culture medium with the drugs listed in Supplemental Table 1. At the end of the treatment the cells were washed in PBS and lysed with SDS denaturing lysis buffer supplemented with Benzonase (4% SDS, 50 mM Tris–HCl, pH 7.4, 100 mM NaCl, 4 mM MgCl_2_, 5 U Benzonase/3 million cells). Cells were lysed directly on the petri dish after carefully removing the PBS and the lysates were collected with the help of a cell scraper. Total protein was determined by NanoDrop (A280 setting) after which the samples were equalized and boiled in Laemmli buffer. Lysates were separated on SDS-PAGE and transferred onto a nitrocellulose membrane. After blocking with 5% non-fat dry milk in TBS with 0.1% Tween 20 the membranes were incubated with primary antibodies, followed by the appropriate HRP-coupled secondary antibodies (Supplemental Table 2). The signal was visualized with commercial ECL HRP substrate (advansta WesternBright or Thermo Scientific Pierce), according to the manufacturer’s instructions, and imaged with a Uvitec Alliance Q9 Advanced system.

### Chromatin fractionation

The chromatin fraction was isolated using the Thermo Scientific Subcellular Protein Fractionation Kit. Briefly, cells were treated for the indicated time points, washed in PBS and then the ice cold cytoplasmic extraction buffer supplemented with protease inhibitor cocktail (PIC), PARGi and Olaparib at a final concentration of 1µM was added. Cells were scraped, collected and incubated on ice for 10 minutes with gentle tapping. Cells were centrifuged at 500 × *g* for 5 minutes at 4 °C. Cytoplasmic fraction was collected in pre-chilled tubes and the pellet was dissolved and shortly vortexed in ice cold membrane extraction buffer supplemented with PIC, PARGi and Olaparib. Tubes were incubated on ice for 10 minutes with gentle tapping and centrifuged at 4000 × *g* for 5 minutes at 4 °C. The pellet was dissolved in ice cold nuclear extraction buffer supplemented with inhibitors, thoroughly vortexed and then incubated for 30 minutes on ice with gentle tapping. Centrifugation was done at 5000 × *g* for 6 minutes at 4 °C. Nuclear fraction was collected in pre-chilled tubes and the pellet was thoroughly vortexed in RT nuclear extraction buffer supplemented with MgCl_2_, micrococcal nuclease, PARGi and Olaparib and incubated for 15 minutes at RT. All fractions were boiled in Laemmli buffer. In addition to the kit-based method, chromatin fractionation was performed by adapting a published protocol (Gillotin, 2018). The final pellet was resuspended in SDS denaturing buffer supplemented with 10 U Benzonase/3million cells, vortexed and incubated at RT until the viscosity was reduced.

### Fluorescent microscopy

After treatment, cells were washed with PBS and fixed in an ice-cold 7: 3 methanol: acetone mixture for 15 minutes at –20 °C. The fixed cells were washed with PBS and permeabilized with 0.2% Triton X-100 in PBS for 10 minutes and then blocked with 5% FBS in PBS for 1 hour at RT. The samples were incubated with indicated primary antibodies followed by fluorescent secondary antibodies diluted in blocking solution (Supplemental Table 2), and finally DAPI was added (1 μg/ml in PBS). Images were acquired using a Zeiss LSM800 confocal microscope equipped with a Plan-Apochromat 20×/0.8 M27 objective and controlled by ZEN 2.3 software. Fluorescence excitation was performed using diode lasers at 405, 488, and 561 nm.

### Flow cytometry

After the indicated treatments, the cells were dissociated with TrypLe Select (Gibco), washed with PBS and fixed with ice-cold ethanol. For labelling the intracellular markers, the cells were permeabilized and blocked with 0.5% Triton X-100 and 5% FBS in PBS, and then incubated with primary antibodies, followed by fluorescently tagged secondary antibodies (Supplemental Table 2). For detecting BrdU incorporation the samples were denatured with 2N HCl and neutralized with citrate-phosphate buffer pH 7.4 prior to adding the anti-BrdU primary antibody. Finally, the DNA staining solution was added (10 μg/ml propidium-iodide and 10 μg/ml RNase in PBS). The samples were analyzed with CytoFLEX S flow cytometer (Beckman Coulter Life Sciences) and the measurement was evaluated with Kaluza Analysis Software (Beckman Coulter Life Sciences).

## Acknowledgements

We thank Adrián Kószó (laboratory of Gyula Timinszky) for technical assistance, and the Flow Cytometry Core Facility at the HUN-REN Biological Research Centre for their support. This work was supported by the National Research Development and Innovation Office (K143248 to G.T. and K142385 to D.S.).

## Author contributions

**Alexandra Mihuț:** Conceptualization, Investigation, Data curation, Formal analysis, Writing – original draft. **Debanjan Ghosh**: Investigation. **Dávid Szüts**: Conceptualization, Writing – review & editing. **Roberta Fajka-Boja**: Conceptualization, Investigation, Data curation, Writing – review & editing. **Gyula Timinszky**: Conceptualization, Supervision, Funding acquisition, Writing – review & editing.

